# Microplate Assay for Denatured Collagen using Collagen Hybridizing Peptides

**DOI:** 10.1101/443242

**Authors:** Allen H. Lin, Jared L. Zitnay, Yang Li, S. Michael Yu, Jeffrey A. Weiss

**Affiliations:** Department of Bioengineering, University of Utah; Scientific Computing and Imaging Institute, University of Utah; Department of Pharmaceutics and Pharmaceutical Chemistry, University of Utah; Department of Orthopaedics, University of Utah

**Keywords:** collagen hybridizing peptide, denatured collagen, microplate assay, osteoarthritis, tendons and ligaments

## Abstract

The purpose of this study was to develop a microplate assay for quantifying denatured collagen by measuring the fluorescence of carboxyfluorescein bound collagen hybridizing peptides (F-CHP). We have shown that F-CHP binds selectively with denatured collagen, and that mechanical overload of tendon fascicles causes collagen denaturation. Proteinase K was used to homogenize tissue samples after F-CHP staining, allowing fluorescence measurement using a microplate reader. We compared our new assay to our previous image analysis method and the trypsin-hydroxyproline assay, which is the only other available method to directly quantify denatured collagen. Relative quantification of denatured collagen was performed in rat tail tendon fascicles subjected to incremental tensile overload, and normal and ostoeoarthritic guinea pig cartilage. In addition, the absolute amount of denatured collagen was determined in rat tail tendon by correlating F-CHP fluorescence with percent denatured collagen as determined by the trypsin-hydroxyproline assay. Rat tail tendon fascicles stretched to low strains (<7.5%) exhibited minimal denature collagen, but values rapidly increased at medium strains (7.5-10.5%) and plateaued at high strains (≥12%). Osteoarthritic cartilage had higher F-CHP fluorescence than healthy cartilage. Both of these outcomes are consistent with previous studies. With the calibration curve, the microplate assay was able to absolutely quantify denatured collagen in mechanically damaged rat tail tendon fascicles as reliably as the trypsin-hydroxyproline assay. Further, we achieved these results more efficiently than current methods in a rapid, high-throughput manner, with multiple types of collagenous tissue while maintaining accuracy.

## INTRODUCTION

Collagen is the most abundant protein in humans and plays a major role in structural support throughout the body. It is involved in essential physiological processes including tissue and extracellular matrix remodeling, injury, and healing.^1^ Damaged or denatured collagen is a major hallmark of many debilitating diseases and injuries, (e.g. cancer, osteoporosis, and osteoarthritis), especially during their progression. For example, it is known that denatured collagen plays a major role in osteoarthritis,^2^ and more recently, has been shown to occur during mechanical damage to tendons.^3,4^ Therefore, the ability to rapidly and accurately quantify denatured collagen would be a powerful asset for the study of many diseases and injuries.

Currently available methods for quantifying denatured collagen are either qualitative, unable to provide information on spatial distribution, or extremely labor intensive. As an example, electron microscopy of mechanically damaged tendons can only reveal qualitative changes in structure, such as the prevalence of kinking and unraveled collagen fibrils.^3^ The trypsin-hydroxyproline (T-H) assay utilizes trypsin, which selectively digests denatured collagen. Here, the hydroxyproline in both denatured and intact collagen are quantified, and the two values used to calculate the percent of collagen that is denatured in the sample. This method has been previously used in both tendons and cartilage.^2,5^ Trypsin is used over other proteolytic enzymes such as pepsin and chymotrypsin due to its specificity for non-triple-helical collagen, while leaving cross-linked telopeptide regions intact.^6^ This prevents the cleavage of hydroxylysine mediated crosslinks which could potentially release triple-helical collagen into the digest, causing overestimation of the amount of denatured collagen. Some investigators have used antibody mediated immunological assays. This was used for articular cartilage, where a monoclonal antibody with affinity for denatured type II and the α3 chain of type XI collagen was utilized.^7^ The methods described above all suffer from various shortcomings. The T-H assay is a time-consuming protocol that spans several days and is prone to error during its many pipetting steps. It is also unable to provide spatial information about the distribution of denatured collagen throughout the samples. The immunological assay only detects specific epitopes of type II collagen and thus it is questionable whether the resulting signal reflect the level of all denatured collagen. In addition, it is limited by the technical challenges and costs associated with antibody based assays, such as non-specific binding.^8^

Recently, collagen hybridizing peptides (CHPs) have been used to identify denatured collagen in mechanically damaged rat tail tendons.^4^ CHPs, also known as collagen mimetic peptides,^9^ have repeated glycine-proline-hydroxyproline (GPO) residues that mimic the native structure of collagen. Historically, CHPs have been used to study the structure and folding of collagen, but more recently have been shown to hybridize unfolded α-strands of all collagen types while showing little affinity to intact triple helical collagen.^10,11^ By conjugating CHPs with fluorescent or bioactive moieties such as biotin, collagen that has become denatured due to injury or disease can be identified and visualized.^4,12^ Previously in our lab, CHPs with a (GPO)_9_ sequence bound to carboxyfluorescein (F-CHP) were used to stain rat tail tendon (RTT) fascicles after they were mechanically loaded. The fascicles were imaged using widefield fluorescence microscopy to measure and quantify fluorescence intensity and by extension collagen damage.^4^ While this method was effective at detecting and localizing denatured collagen in a tissue sample, it is unable to quantify the exact amount of denatured collagen, which is important for elucidating trends and mechanisms during disease onset and progression.

Given the shortcomings of previously described methods and the promising results obtained recently using CHPs, we sought to develop a rapid, high throughput method of quantifying denatured collagen based on CHP binding that could be employed using common laboratory equipment. Thus, we utilized F-CHPs in a microplate format. By staining collagenous tissues containing denatured collagen with F-CHP and then homogenizing the tissue, we were able to quantify CHP fluorescence with a microplate reader - and correlate this fluorescence with the amount of denatured collagen. We demonstrated the versatility of this new assay by detecting denatured collagen in both mechanically damaged tendon and osteoarthritic articular cartilage and showed that the assay could be combined with data from the T-H method for absolute quantification of denatured collagen in tendon. Using our method, denatured collagen can be quantified quickly and accurately to give insight to the role of denatured collagen due to injury, disease, growth, and remodeling.

## METHODS

### Experimental Design

For relative quantification, we prepared mechanically damaged rat tail tendon (RTT) fascicles.4 Fascicles were loaded in uniaxial extension to 5, 7.5, 9, 12, and 13.5% strain (n=3 each) and then stained with F-CHP. Unloaded samples (n=3) served as a negative control. To reproduce the procedure used for relative quantification in our previous study, RTT fascicles were imaged using widefield fluorescence microscopy and the mean pixel intensity at each strain level was measured. The clamped regions of the same samples were then trimmed off, leaving just the loaded portion. These samples were weighed and the wet weights were recorded before undergoing the CHP microplate assay described below to quantify fluorescence (Fig. 1).

**Figure 1.**
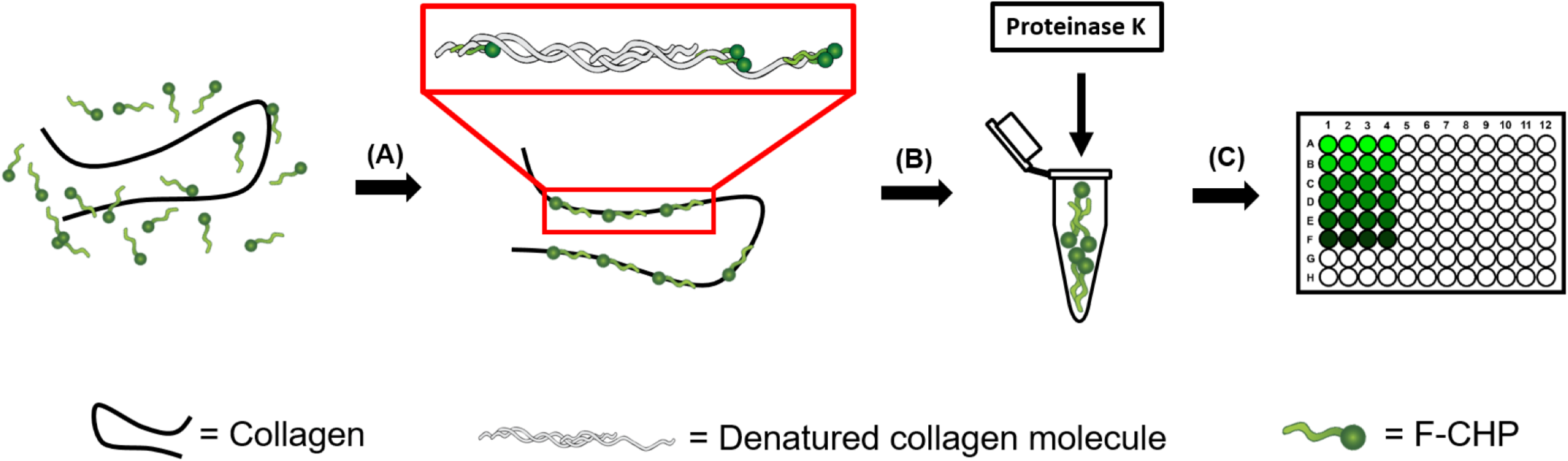
(page width). Schematic illustrating the steps in the CHP microplate assay. (A) After incubation in F-CHP, the sample is rinsed, removing unbound F-CHP. (B) The sample is digested with proteinase K to homogenize it. (C) The fluorescence of the digest is quantified on a 96-well microplate reader.

To achieve relative quantification of denatured collagen in articular cartilage, both young (3-6 months) and old (9-12 months) guinea pig legs were acquired (BioreclamationIVT, Westbury, NY) to compare denatured collagen amounts in healthy and osteoarthritic cartilage. Guinea pigs exhibit osteoarthritis in the large joints upon reaching skeletal maturity, allowing comparisons between young (healthy) and old (osteoarthritic) cartilage.^13^ Two sets of 4 mm plugs (n=5 per age group per set) were taken from the center of the medial tibial plateau using a trephine drill bit (Fig. 2). The samples were frozen in 1xPBS at -80°C until they were used. The plugs were embedded in optimal cutting temperature compound and the trabecular bone was removed using a sledge microtome, leaving cartilage disks. One set underwent trypsin digestion, where the hydroxyproline in the digest was quantified and normalized to wet weight of tissue. The other set of cartilage disks underwent the CHP assay, where denatured collagen was quantified by fluorescence. When we performed the CHP assay on the cartilage disks, calcified cartilage remained. The remaining tissue was hydrolyzed with HCl and the hydroxyproline content measured to determine if collagen remained within the calcified cartilage (see Supplementary Methods).

**Figure 2.**
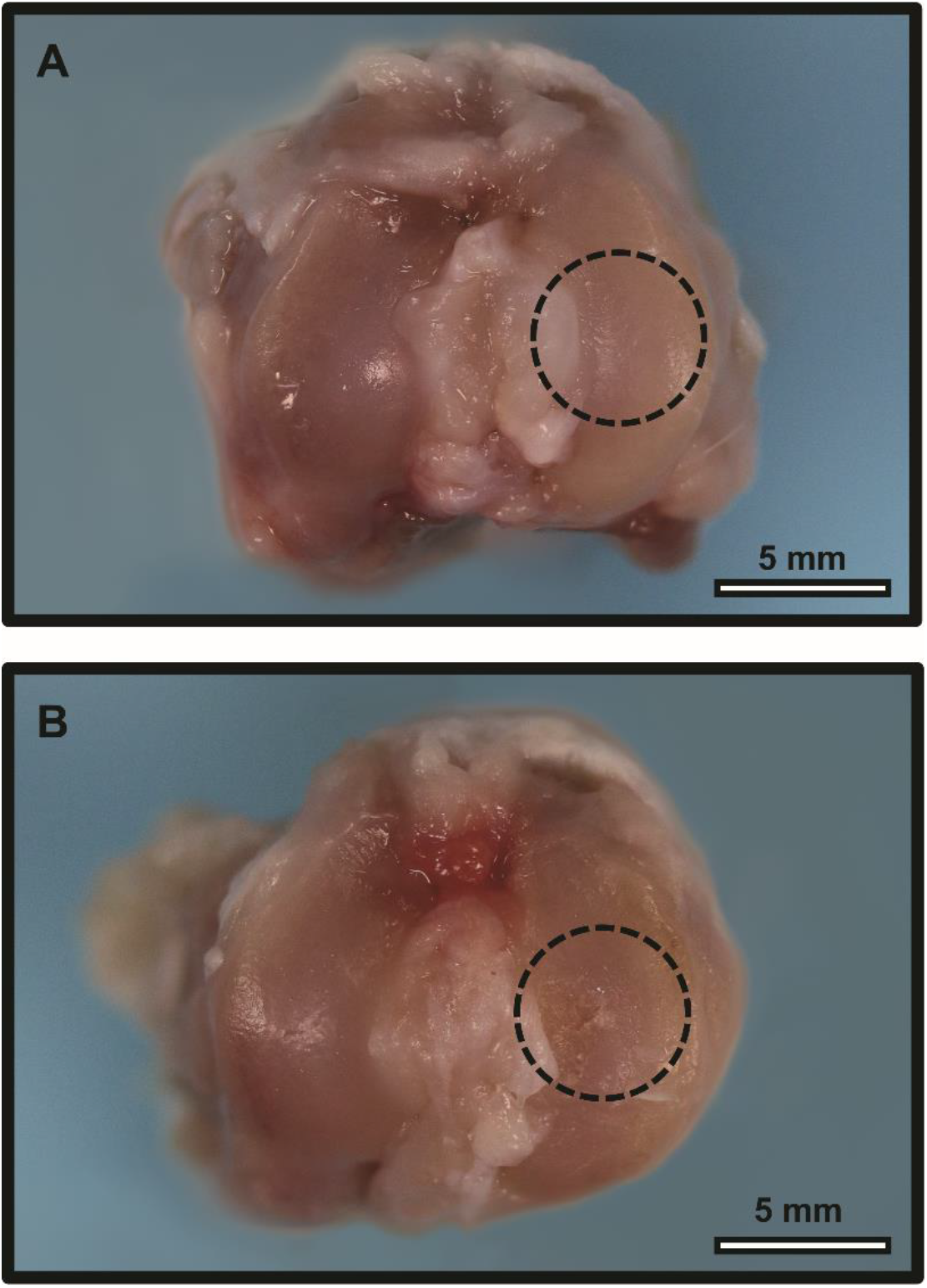
(column width). Representative images of cartilage from the left knee of (A) young (healthy) and (B) old (osteoarthritic) guinea pigs. Dashed circles indicate the regions of the medial tibial plateaus from which samples were taken.

A calibration curve was developed to relate denatured collagen to CHP fluorescence. To create graded amounts of denatured collagen, RTT fascicles were subjected to controlled application of heat. Two sets of RTT fascicles were heated at 57°C for 0, 3, 5, 7, or 9 minutes (n=5 per set). One set of samples was used for the T-H assay, while the other set was used for the CHP microplate assay. The results of the two assays were correlated to create a calibration curve, giving the ability to report the percent denatured collagen based on CHP fluorescence readings.

The accuracy of the calibration curve was tested by performing the T-H and CHP microplate assays on two additional sets of mechanically damaged RTT fascicles. RTT fascicles were stretched to strains of 0, 5, 9, and 12% (n=3 per set). The calibration curve correlated the fluorescence with percent denatured collagen, which was compared to the value calculated from the T-H assay.

### Mechanical Damage to Tendon Fascicles

The approach for creating mechanical damage to tendon fascicles followed the protocol in our previous study.^4^ Rat tail tendon fascicles were dissected from 8 week old Sprague Dawley rat tails (BioreclamationIVT, Westbury, NY) by twisting the distal end of the tail with a pair of hemostats until the skin separated and pulling the distal end away, revealing the fascicles. Fascicles were mechanically damaged by applying a prescribed amount of uniaxial tensile strain on a material testing machine (Electroforce 3330-AT Series II, TA Instruments, Eden Prairie, MN). A 0.03 N preload was applied, and the fascicles were strained at a rate of 0.5% s^-1^ at room temperature in 1xPBS. Upon reaching the prescribed strain level, samples were immediately unloaded. The clamped ends of the samples were trimmed off to isolate the loaded region of the fascicle, which was stored in 1.0 mL of 1xPBS at 4°C until CHP staining.

### CHP Staining

Each sample was weighed and the wet mass was recorded. A stock solution of 150 μM F-CHP (3Helix, Inc., Salt Lake City, UT) was heated at 80°C for ten minutes and then rapidly cooled with 4°C water for 5 seconds to reverse the CHP trimerization that naturally occurs due to its collagen mimetic structure.^10^ Samples were placed in 450 μL of 1xPBS, to which 50 μL of 150 μM F-CHP was added for a final concentration of 15 μM. The samples were stained for at least 12 hours at 4°C with gentle agitation. Afterwards, the samples were rinsed with 1.0 mL of 1xPBS at room temperature with gentle agitation for at least 30 minutes. RTT fascicles and cartilage were rinsed three and five times, respectively, to remove excess unbound F-CHP. Samples were stored at -20°C in 1.0 mL of 1xPBS until used for either the microplate assay or widefield fluorescence imaging.

### Widefield Fluorescence Microscopy

RTT fascicles stained with F-CHP were placed between a glass slide and cover slip with 1xPBS and imaged at 4x magnification using the fluorescein isothiocyanate (FITC) channel with a CCD camera at 16-bit (Ti-E, Nikon, Tokyo, Japan). Captured images were thresholded using Otsu’s method^14^ to isolate the stained fascicle region, and the mean pixel intensity value was calculated, as per our previous method.^4^ Image processing was performed using MATLAB (2016b, MathWorks, Natick, MA).

### CHP Microplate Assay

To enable the detection of F-CHP fluorescence using a microplate reader, samples were homogenized in 500 μL of a 1 mg/mL proteinase K (Cat No. 193981, MP Biomedicals, LLC, Santa Ana, CA) solution in deionized water for three hours at 60°C. The amount of time required for digestion was determined by digesting samples for different periods of time, and observing when the fluorescence reached a plateau. After homogenization, the fluorescence of 200 μL duplicates was measured at excitation and emission wavelengths of 485 and 525 nm, respectively on a microplate reader (SpectraMax Gemini XPS, Molecular Devices, Sunnyvale, CA). The mean value of duplicate fluorescence measurements was then normalized by the wet weight of the sample.

### Trypsin-Hydroxyproline (T-H) Assay

The T-H assay was performed on both RTT fascicles and guinea pig cartilage samples (see Supplementary Methods). For RTT fascicles, the percent collagen digested by trypsin was calculated using equation S-1. The cartilage samples contained both articular and calcified cartilage, the latter of which trypsin was unable to completely diffuse through. This caused an underestimation of the percent collagen denatured in the sample. Therefore, the amount of collagen in articular cartilage digested by trypsin was normalized to the sample wet weight for relative quantification between healthy and osteoarthritic cartilage.

### Statistical Analysis

For both the widefield fluorescence imaging and CHP microplate assay, the mean value of fluorescence intensity for each strain level was compared to unloaded controls using a one-tailed t-test with significance established at the α=0.05 level. The amount of denatured collagen measured in healthy and osteoarthritic cartilage was compared a two-tailed t-test (α=0.05) for both T-H and microplate assays. The calibration curve correlating normalized fluorescence with percent collagen denatured was fitted with a simple linear regression. Bland-Altman^15^ analysis was used to compare the percent denatured collagen determined by the T-H and CHP microplate assays with 95% limits of agreement. The two absolute quantification methods used for RTT fascicles were also compared at each strain group using a two-tailed t-test (α=0.05) to determine if the result from the methods were significantly different. Statistical analyses were performed with SigmaPlot 13.0 (Systat Software, Inc., San Jose, CA).

## RESULTS

### Widefield Fluorescence Microscopy vs. CHP Microplate Assay

The fluorescence of mechanically damaged RTT fascicles stained with F-CHP as a function of applied strain exhibited the same trend when quantified with both widefield fluorescence microscopy and the CHP microplate assay (Fig. 3). Both revealed that above a strain threshold, the F-CHP intensity and by extension the extent of damage to the collagen molecules in the fascicle increased with strain before it plateaued, exhibiting a sigmoidal shape. When fascicles were stretched up to 7.5% strain, there was minimal fluorescence indicating that single loading events at these strains do not cause permanent denaturation. At strains beyond 7.5%, the mechanical damage caused irreversible denaturation of the collagen molecule to rapidly accumulate, before plateauing at strains above 12%. For both widefield fluorescence microscopy and the CHP microplate assay, the mean fluorescence measured for all strain groups at or above 7.5% was significantly greater than the unloaded controls (p<0.05). The fluorescence of the 5% strain group was significantly greater than unloaded controls for only widefield fluorescence microscopy (p=0.0249). The two methods provided similar precision, as evidenced by the error bars.

**Figure 3.**
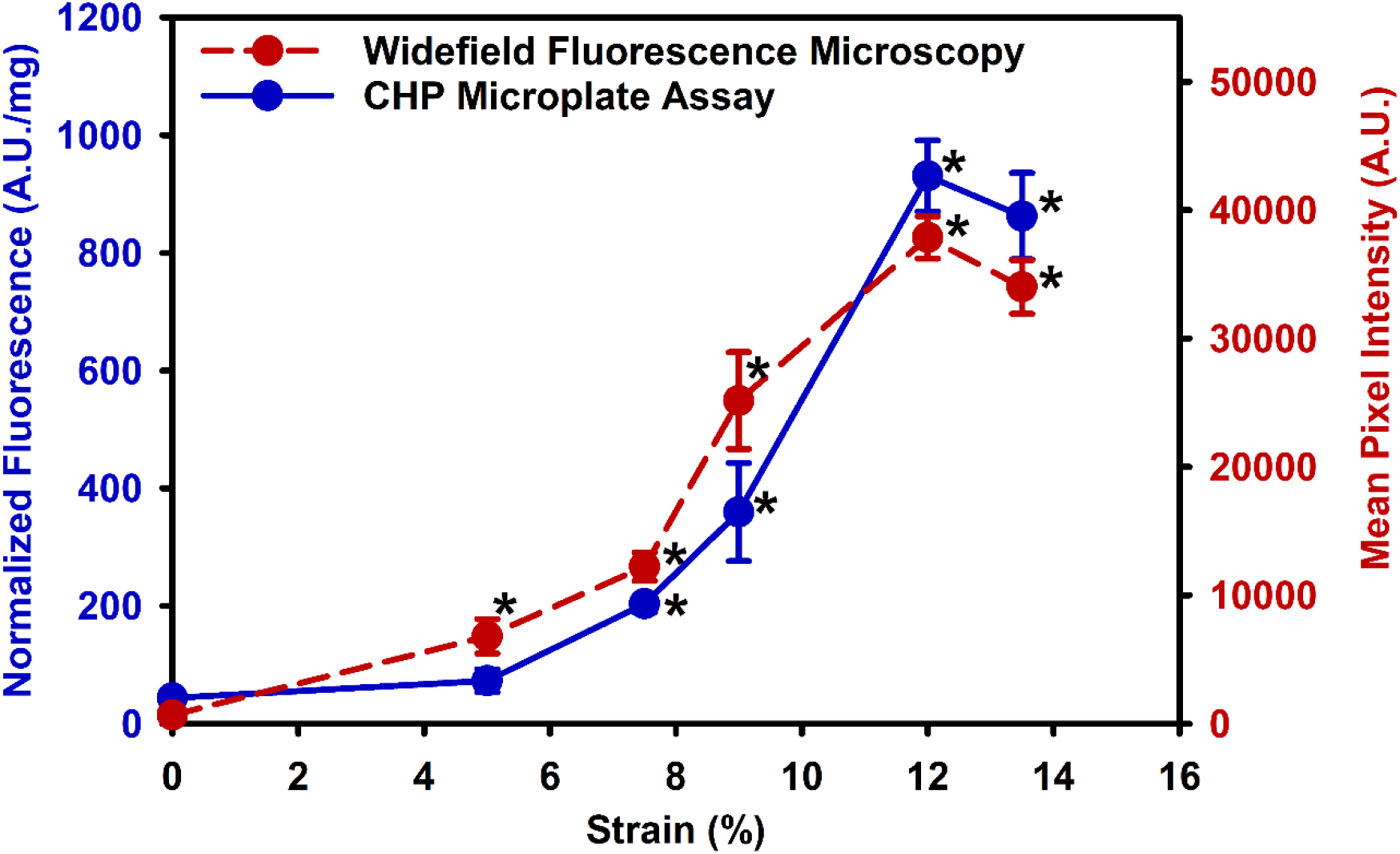
Normalized F-CHP fluorescence as measured by the CHP microplate assay (blue) and mean pixel intensity as measured by widefield fluorescence microscopy (red) for F-CHP stained RTT fascicles. Both methods exhibited the same trend in fluorescence relative to strain. The low fluorescence intensity at 0% strain originate from baseline levels of denatured collagen due to collagen remodeling which occurs in healthy tissues. At intermediate (7.5-10.5%) strains, collagen denaturation from mechanical damage begins to accumulate, as evident by the increased fluorescence from more F-CHP binding sites. At high (>12%) strains, the magnitude of collagen denaturation plateaus. Mean±StDev. *Statistically different from the unloaded control group at p<0.05.

### Calibration Curve for Direct Quantification in RTT Fascicles using CHP Microplate Assay

By controlled heating of RTT fascicles for varying times, we created graded amounts of denatured collagen that could be distinguished by both the CHP microplate and T-H assays (Fig. 4). Greater amounts of denatured collagen was observed for longer heating times, both in terms of CHP binding and fluorescence and the T-H assay. The calibration curve relating normalized F-CHP fluorescence and percent denatured collagen was fit using linear regression, with R^2^=0.900.

**Figure 4.**
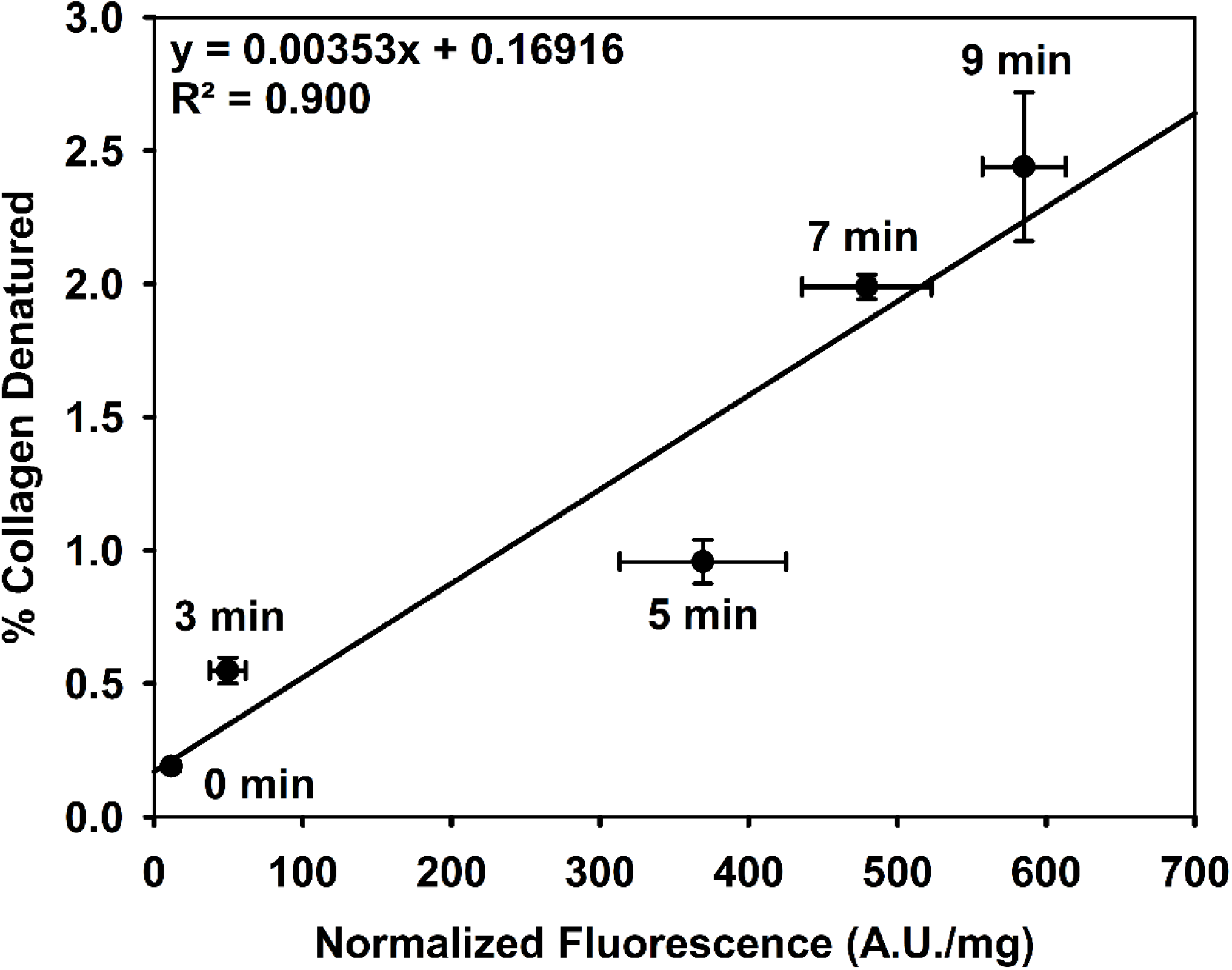
Calibration curve for RTT fascicles correlating normalized fluorescence from F-CHP with percent denatured collagen from the T-H assay. The amount of denatured collagen in RTT fascicles increased linearly with heat denaturation time (indicated at each data point) as measured by both methods, and the measurements from both methods were strongly correlated (R^2^ = 0.900). Error bars represent 95% confidence intervals.

### Comparison of T-H and CHP Microplate Assays for RTT Fascicles

Both the T-H and CHP microplate assays were performed on mechanically damaged RTT fascicles. The results of the CHP microplate assay were converted to percent denatured collagen using the established calibration curve and compared to the T-H assay (Fig. 5, top). The two methods showed very good agreement, with no significant differences found within all four strain groups at α=0.05. Bland-Altman analysis comparing the CHP microplate and T-H assays revealed a bias of -0.1028% with 95% limits of agreement being [-0.3756%, 0.1699%] (Fig. 5, bottom). The low bias indicated very good agreement between the two methods of quantifying denatured collagen.

**Figure 5.**
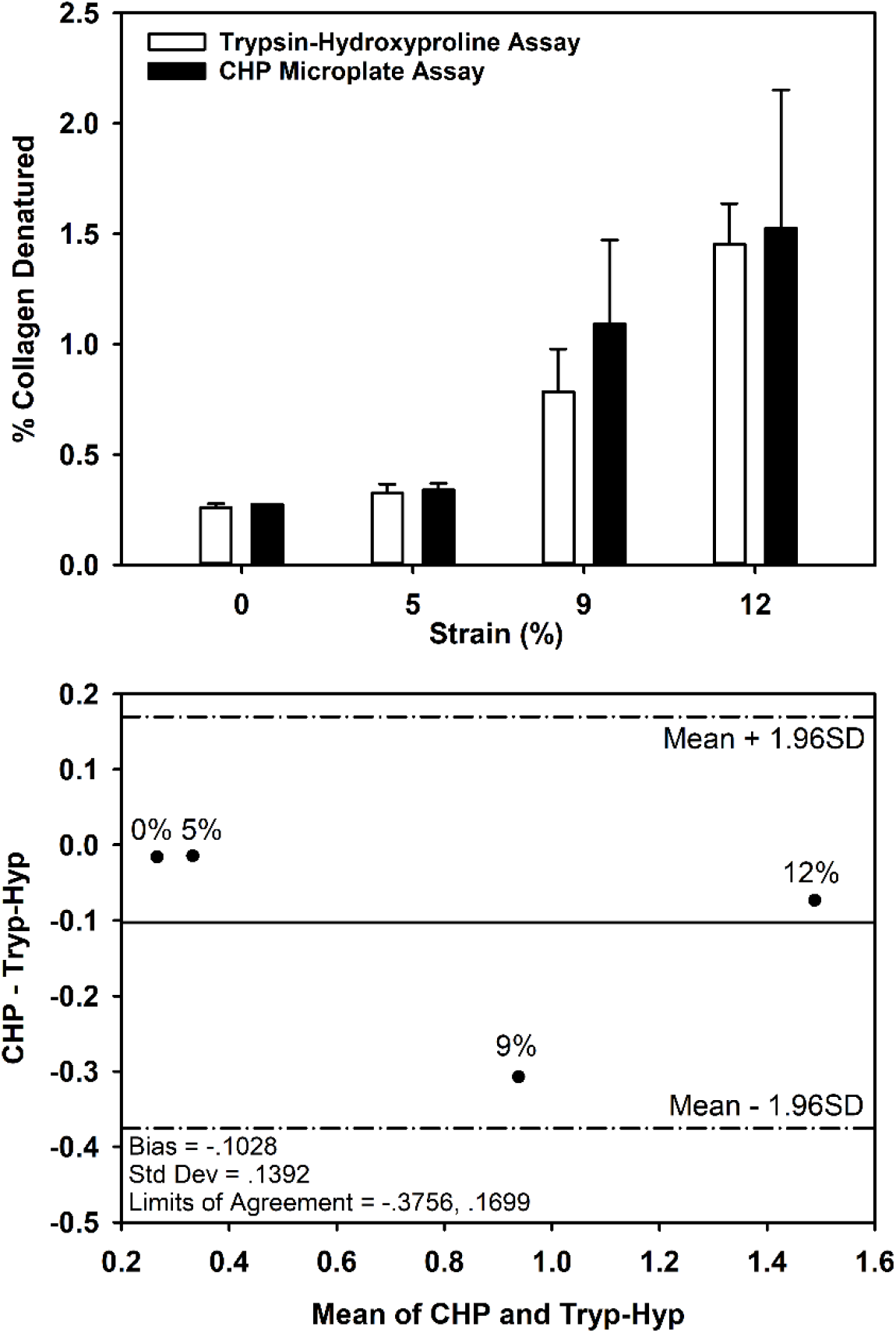
(column width). Top: Percent denatured collagen in mechanically damaged rat tail tendon fascicles as determined by the CHP microplate (closed bars) and T-H (open bars) assays. No significant difference was found between mean values (α=0.05) within all four strain groups. Mean±S.E.M. Bottom: Bland-Altman plot with the difference and mean values between the CHP and T-H assays for rat tail tendon fascicles stretched to several strains. A mean bias of - 0.1028 and 95% limits of agreement of [-0.3756, 0.1699] indicate good agreement between the two assays.

### Comparison of T-H and CHP Microplate Assays for Cartilage

Denatured collagen in healthy and young guinea pig cartilage was quantified by both the CHP assay and the T-H assay (Fig. 6). The young guinea pig cartilage appeared healthy and normal while the old cartilage exhibited signs of osteoarthritis such as fibrillation on the surface of the medial tibial plateau (Fig. 2).^2^ Both methods measured a higher level of denatured collagen in osteoarthritic cartilage than healthy cartilage. The results of the CHP assay demonstrated a significantly greater amount of denatured collagen in the OA cartilage than the healthy cartilage (p<0.05). Interestingly, there was no statistical difference in the amount of denatured collagen between normal and OA cartilage as measured by the T-H assay (p=0.249).

**Figure 6.**
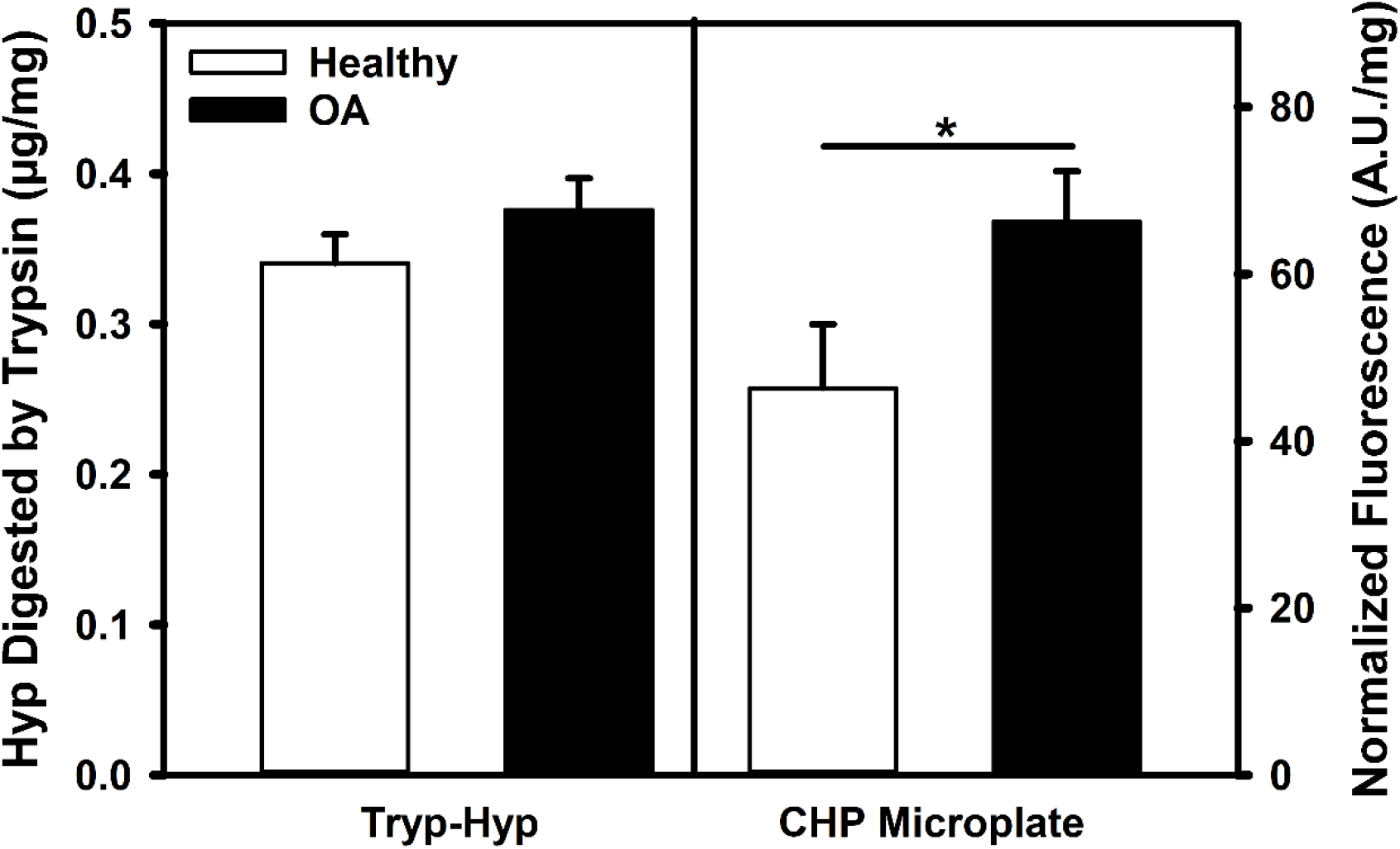
Greater levels of denatured collagen were observed in the osteoarthritic guinea pig cartilage relative to the healthy cartilage, for both trypsin digestion and the CHP assay. Mean±S.E.M. There was a significant difference between groups for the CHP assay, but not the trypsin digestion. *P<0.05.

## DISCUSSION

We developed and verified a rapid, high throughput assay to determine the amount of denatured collagen in tissue samples based on quantifying fluorescence produced by F-CHP bound to denatured collagen with a microplate reader. The assay is based on the fact that CHP remains bound to denatured collagen even after standard tissue digestion protocols are used. This new method absolutely quantified denatured collagen as reliably yet more efficiently than the T-H assay, requiring less than half the time necessary while also entailing fewer steps and supplies. It is also as accurate as our previous method for measuring CHP fluorescence using widefield fluorescence microscopy, but it is not subject to the same variability that can result from variations in imaging parameters and image processing. Absolute quantification of denatured collagen initially requires the creation of calibration curves correlating normalized F-CHP fluorescence with percent denatured collagen for each specific tissue type. However, once a calibration curve is established, the amount of denatured collagen in tissue samples can be quantified from F-CHP fluorescence without the need to perform further calibration assays. There was no statistical difference between the two methods when this approach was used for absolute quantification of denatured collagen in damaged RTT fascicles, demonstrating its efficacy. Bland-Altman analysis supported this, as evidenced by the low bias of -0.1028%, which was mostly influenced by the 9% strain group. The narrow range for the 95% limits of agreements indicated that most differences in outcomes between the two assays should be minimal.

The CHP microplate assay produced the same trend for fluorescence as a function of applied strain in RTT fascicles as widefield fluorescence microscopy. F-CHP fluorescence quantification by widefield fluorescence microscopy is limited to two-dimensional tissue samples, prone to image processing errors, and time consuming as each sample must be individually placed on a slide and imaged. This contrasts with the CHP microplate assay, where the samples are homogenized with proteinase K, allowing simultaneous quantification of samples using the 96-well plate format. By digesting the F-CHP bound tissue into a homogenous solution, this assay also in theory enables more precise measurements of fluorescence intensity. Minimal CHP binding was observed at low levels of strain (<7.5%), indicating a lack of permanent denaturation of collagen under these sub-failure loading conditions. The small amount of fluorescence observed at these lower strain levels was likely due to denatured collagen from collagen remodeling that naturally occurs in healthy tissues. At medium strains (7.5-10.5%), permanent collagen denaturation occurred, as evidenced by a sharp increase in fluorescence from increased CHP binding. At high strains, the fluorescence intensity plateaued, demonstrating that there is no longer an increase in binding sites for CHPs. These observations were seen using both F-CHP fluorescence quantification techniques, showing that the CHP microplate assay reliably measures relative amounts of F-CHP, to the same extent as widefield fluorescence microscopy, while doing so more efficiently.

The mean fluorescence of the 5% strain samples was significantly greater than unloaded controls when quantified by widefield microscopy. This was unexpected, for significance was not observed when fluorescence was quantified by the microplate assay. A possible explanation for this difference is that the image analysis technique for widefield microscopy is subject to error, especially for low strain samples. The entire sample was imaged during widefield microscopy, while the unclamped portions of the samples were removed for the CHP assay. Therefore, CHP binding to untested regions of the sample could have caused over-reporting of the fluorescence intensity, which would be more prevalent at low strain samples. Since the unloaded regions were trimmed off for the CHP microplate assay, the fluorescence measured by our new method was only representative of the collagen denatured in the loaded region.

We also demonstrated relative quantification of denatured collagen in cartilage. Our data supports previous reports of elevated levels of denatured collagen occurring in osteoarthritic cartilage.^2,7^ We observed greater F-CHP fluorescence from the CHP assay, and greater levels of hydroxyproline in the supernatant following trypsin digestion in osteoarthritic cartilage. When the cartilage disks were digested with proteinase K, leftover tissue was observed, later determined to be calcified cartilage.^16^ The normalized amount of hydroxyproline measured in the leftover tissue was comparable to that of the original sample, indicating that proteinase K was unable to digest collagen in the calcified tissue. Due to proteinase K and trypsin sharing similar molecular weights (28.9 kDa vs 24 kDa),^17,18^ it is likely that trypsin also failed to diffuse into the calcified portion and digest denatured collagen. Previous studies have also reported reduced diffusivity in calcified cartilage compared to non-calcified cartilage.^19^ Hence, the F-CHP fluorescence and hydroxyproline quantified following trypsin digestion both originate from non-calcified cartilage, still allowing relative comparisons between the two methods.

The relative amount of denatured collagen in osteoarthritic cartilage was significantly greater for the CHP assay, but not trypsin digestion. This could be due to normalizing the results of both assays to the wet weight of the entire sample. Different samples could have contained different relative amounts of calcified and non-calcified cartilage depending on how much calcified cartilage was removed during sample preparation. Samples with greater calcified cartilage fractions would result in underestimation of the amount of denatured collagen. We were unable to quantify the wet weight or hydroxyproline amount of the calcified cartilage since the T-H assay is completely destructive. We did not observe calcified cartilage in pilot studies that we performed using thicker cartilage samples originating from bovine or porcine sources due to the ease of removing the calcified region. For these tissues, the proteinase K incubation resulted in complete digestion of these samples.

We bridged the gap between two separate methods of quantifying denatured collagen – the T-H assay and CHP fluorescence – by establishing a calibration curve correlating them. The calibration curve exhibited a linear trend with R^2^=0.900. The mean of 5-minute group was furthest form the regression line, and could be due to sample-to-sample variability or slight perturbations in temperature. Around 57.0°C, variations as little as 0.5°C can cause significant changes in the thermal denaturation of collagen.^20^ This curve reliably determined absolute amounts of denatured collagen from CHP fluorescence, which alone can only measure relative amounts. The T-H method requires over 64 hours of assay time,^4^ and is prone to error with its complicated multistep protocol. The CHP microplate assay contrasts with this by requiring less than 24 hours of bench time following creation of the calibration curve, requiring only a simple staining process followed by tissue digestion. This protocol will allow researchers to rapidly quantify the percentage of collagen denatured in connective tissues. Given the prevalence and role of denatured collagen in injuries and pathologies such as osteoarthritis, cancer, and fibrosis,^12^ the method can help provide insight on the onset and progression of these ailments.

The other method of absolutely quantifying denatured collagen, an immunoassay, has seen use in cartilage. Denatured collagen is detected using a monoclonal antibody that reacts with an epitope found in denatured but not intact type II collagen using an enzyme-linked immunosorbent assay. However, it is unclear if these epitopes are exposed prior to catastrophic failure of the collagen molecule. In the past, this method was used to identify collagen damage following indentation of osteochondral plugs.^21,22^ Depending on the load applied, the Col2-3/4m antibody was unable to detect the exposed epitope and denatured collagen.^23^ Healthy unloaded cartilage stained with F-CHP have revealed the presence of denatured collagen.^12^ This suggests that CHPs are more sensitive for detecting denatured collagen compared to immunological methods, and are able to detect denatured collagen prior to degradation of the collagen matrix.^24^ Overall, the CHP microplate assay is more versatile than existing methods of absolutely quantifying denatured collagen, while being more efficient.

The collagen mimetic motif in CHP theoretically allows it to bind with denatured collagen of any type, and its use has been demonstrated in tissues containing collagen types I, II, III, and IV.^12^ Therefore, this assay could potentially be used with any type of collagenous tissue. When applied to mechanically damaged RTT fascicles of varying strains, no statistical difference was found between the values calculated from the calibration curve and the T-H assay (Fig. 5). The greatest bias was seen with the 9% strain group. This is likely due to the fact that 9% strain was in the region of strain where the samples departed linear stress-strain behavior which is associated with permanent tissue damage.^4^ Therefore, sample-to-sample variation alone could result in a relatively large changes in the amount of denatured collagen compared to low strains and the plateau region at high strains. In addition, the amount of denatured collagen was similar to that found by Zitnay et al^4^ for the same levels of damage. The calibration curve reliably converted fluorescence to percent collagen denatured despite being created with heat denatured collagenous tissue samples, demonstrating that the binding mechanism of CHPs to denatured collagen is the same regardless of denaturation method.

Several points need to be considered when applying the CHP microplate assay. Due to their collagen mimetic structure, CHPs will self-assemble into trimers over time, photo-quenching the fluorophore and preventing binding to denatured collagen.^10^ This issue was resolved in this study by heating the F-CHP solution prior to tissue staining to melt the CHP trimers. In addition, the diffusion characteristics of CHPs in collagenous tissues have not been characterized, so the optimal staining time for CHP penetration and washing time must be determined by iterative experiments for each new tissue type and for significant changes in tissue geometry. Additionally, it is unknown if the calibration curves remain linear beyond the levels of denatured collagen observed in this study. However, relative quantification of denatured collagen should always be possible.

## ACKNOWLEDGMENTS

Financial support from NIH #R01EB015133 and R01AR071358 is gratefully acknowledged. Imaging was performed at the Fluorescence Microscopy Core Facility, a part of the Health Sciences Cores at the University of Utah. Microscopy equipment was obtained using a NCRR Shared Equipment Grant # 1S10RR024761-01. Y.L. and S.M.Y. are co-founders of 3Helix, Inc. which commercializes the collagen hybridizing peptides. All other authors have no professional or financial conflicts of interest to disclose.

AUTHORS’ CONTRIBUTIONS
A.H.L., J.L.Z., Y.L., S.M.Y., and J.A.W. designed the study; A.H.L., J.L.Z., and Y.L. performed the experiments; A.H.L., J.L.Z., Y.L., S.M.Y., and J.A.W. analyzed the results. All authors contributed to writing and editing the manuscript, and approve of the final manuscript.

